# Mitochondrial DNA and their nuclear copies in parasitic wasp *Pteromalus puparum*: A comparative analysis in Chalcidoidea

**DOI:** 10.1101/396911

**Authors:** Zhichao Yan, Qi Fang, Yu Tian, Fang Wang, Xuexin Chen, John H. Werren, Gongyin Ye

**Affiliations:** State Key Laboratory of Rice Biology & Ministry of Agriculture Key Laboratory of Molecular Biology of Crop Pathogens and Insects, Institute of Insect Sciences, Zhejiang University, Hangzhou 310058, China.; Department of Biology, University of Rochester, Rochester, NY 14627, USA.; Nextomics Biosciences Co., Ltd., Wuhan, Hubei 430073, China.

**Author notes:** To whom correspondence should be addressed: Ye Gongyin, Institute of Insect Sciences, C1152 Agro-Bio Complex, Zhejiang University, Hangzhou 310058, China. John H. Werren, Department of Biology, University of Rochester, Rochester, NY 14627, USA.

**Keywords:** mitochondrial genome, NUMTs, gene order rearrangement, evolutionary rate, Chalcidoidea

## Abstract

Chalcidoidea (chalcidoid wasps) are an abundant and megadiverse insect group with both ecological and economical importance. Here we report a complete mitochondrial genome in Chalcidoidea from *Pteromalus puparum* (Pteromalidae). Eight tandem repeats followed by 6 reversed repeats were detected in its 3,308 bp control region. This long and complex control region may explain failures of amplifying and sequencing of complete mitochondrial genomes in some chalcidoids. In addition to 37 typical mitochondrial genes, an extra identical isoleucine tRNA (trnI) was detected. We speculate this recent mitochondrial gene duplication indicates that gene arrangements in chalcidoids are ongoing. A comparison among available chalcidoid mitochondrial genomes, reveals rapid gene order rearrangements overall, and high substitution rate in *P. puparum*. In addition, we identified 24 nuclear sequences of mitochondrial origin (**NUMTs**) in *P. puparum*, summing up to 9,989 bp, with 3,617 bp of these NUMTs originating from mitochondrial coding regions. NUMTs abundance in *P. puparum* is only one-twelfth of that in its relative, *Nasonia vitripennis*. Based on phylogenetic analysis, we provide evidence that a faster nuclear degradation rate contributes to the reduced NUMT numbers in *P. puparum*. Overall, our study shows unusually high rates of mitochondrial evolution and considerable variation in NUMT accumulation in Chalcidoidea.

## Introduction

Chalcidoidea in Hymenoptera is one of the most megadiverse groups of insects, and is both ecologically and economically important (Heraty *et al.,* 2013; Peters *et al.,* 2018). With an estimation of more than 500,000 species (Peters *et al.,* 2018), Chalcidoidea has an astonishing diversity of morphology and biology, and is present in virtually all extant terrestrial habitats (Gibson *et al.,* 1999; Noyes 2016). Most chalcidoids are parasitoids, while some are phytophagous. Parasitic chalcidoids are widely and successfully used as biological control agents for various agricultural pests and arthropod vectors of human disease (Heraty 2017).

Little information is known about mitochondrial genomes and their evolution in Chalcidoidea. Only a few mitochondrial genomes have been reported in this group (Chen *et al.,* 2018; Nedoluzhko *et al.,* 2016; Oliveira *et al.,* 2008; Su *et al.,* 2016; Xiao *et al.,* 2011; Xiao *et al.,* 2012; Yang *et al.,* 2018; Zhu *et al.,* 2018). Mitochondrial control regions were reported difficult to be amplified in Chalcidoidea for unknown reasons (Oliveira *et al.,* 2008; Xiao *et al.,* 2011; Xiao *et al.,* 2012; Zhu *et al.,* 2018). In *Nasonia*, several PCR conditions using all possible combinations failed in joining two mitochondrial coding fragments (Oliveira *et al.,* 2008). Although sampling is limited, studies revealed rapid evolution of mitochondrial genomes in Chalcidoidea. Dramatic gene order rearrangement and extremely high substitution rates are observed (Chen *et* al., 2018; Oliveira *et al.,* 2008; Xiao *et al.,* 2011; Xiao *et al.,* 2012; Yang *et al.,* 2018). A striking example is that intraspecific synonymous substitution rates in mitochondria between *Wolbachia* infected and uninfected fig wasp *Ceratosolen solmsi* (Agaonidae) is even higher than the mitochondrial interspecific synonymous substitution rates among *Drosophila* species (Xiao *et al.,* 2012). In addition, Raychoudhury *et al.* (2009b) found in the Nasonia species complex that mitochondrial synonymous substitution rates are 40-fold greater than nuclear rates, and 32-fold greater than their associated *Wolbachia*. These findings suggest a highly elevated mitochondrial mutation rate in chalcidoid wasps. Extremely rapid evolution makes Chalcidoidea an intriguing group for studying mitochondrial evolution, but also introduce difficulties especially for long-PCR-based mitochondrial sequencing (Burger *et al.,* 2007). This may have contributed to poor sampling of mitochondrial genomes in the group.

The mitochondrion is the product of an ancient bacterial symbiosis (Pittis & Gabaldon 2016; Sagan 1967). Most functional mitochondrial genes have been lost or transferred into nuclear genomes (Rand *et al.,* 2004). In plants, the transferring of functional mitochondrial genes into nuclei is still an ongoing process (Bensasson *et al.,* 2001; Henze & Martin 2001). By contrast, animal mitochondrial genomes now are compact with 37 genes, and there are no reports of recent transfer of mitochondrial genes to the nucleus with subsequent loss in the mitochondria (Bensasson *et al.,* 2001; Rand *et al.,* 2004). However, the transferring of mitochondrial DNA into nuclear genomes is still an ongoing process in both plants and animals (Bensasson *et al.,* 2001; Hazkani-Covo & Covo 2008; Hazkani-Covo *et al.,* 2010; Tsuji *et al.,* 2012). In addition, nuclear sequences of mitochondrial origin (**NUMTs**) are prevalent in animal and plant genomes, and are proposed as a window for studying how mitochondrial genes are transferred into nuclear genomes (Hazkani-Covo & Covo 2008; Tsuji *et al.,* 2012). NUMTs are also good materials for studying nuclear sequence evolution without selective constraints (Bensasson *et al.,* 2001). NUMTs have been investigated in several insect genomes, revealing a great variation in abundance, from as much as over 230 kb in *Apis mellifera* genome (246.9 Mb) to none in *Anopheles gambiae* genome (250.7 Mb) (Behura 2007; Hazkani-Covo et al., 2010; Richly & Leister 2004; Viljakainen et al., 2010). The reasons for interspecific variation of NUMTs are still poorly understood. Although a correlation was reported between NUMTs abundance and nuclear genome size, it poorly explained the dramatic variation of NUMTs abundance, especially among species with similar genome sizes (Hazkani-Covo *et al.,* 2010; Song *et al.,* 2013). Other factors were also proposed to explain the interspecific diversity of NUMTs abundance, such as the mitochondrial genome structure (linear or circular) (Song *et al.,* 2013), number of mitochondria in the germline (Smith *et al.,* 2011), or species-specific mechanisms controlling deletion of nuclear DNA (Hazkani-Covo *et al.,* 2010; Richly & Leister 2004).

Here we report a complete mitochondrial genome and their NUMTs in the endoparasitoid *Pteromalus puparum* (Chalcidoidea: Pteromalidae). Comparative analyses were also conducted to other Chalcidoidea. In summary, we 1) assembly the *P*. *puparum* circular mitochondrial genome including control region, 2) show frequent gene order rearrangement in Chalcidoidea, 3) find high protein substitution rate with considerable variation in Chalcidoidea mitochondrial genomes, and 4) detect fewer NUMTs in *P. puparum* than its relative, *N. vitripennis*.

## Results and Discussion

### 1, *Pteromalus puparum* mitochondrial genome

A complete *P. puparum* mitochondrial genome was retrieved from the genome sequencing project of *P. puparum* (see methods). Combined with both Illumina and PacBio reads, we were able to assemble a circular *P. puparum* mitochondrial genome (Figure 1). It is the first complete mitochondrial genome for the family Pteromalidae to our knowledge, although there are partial mitochondrial genomes available for *Nasonia* and *Philotrypesis* (Oliveira *et al.,* 2008; Xiao *et al.,* 2011). As shown in Figure S1 and Table S1, 92 PacBio reads spanned the control region with good coverage, supporting the joining between the coding region and the control region. The total length of circular mitochondrion is 18,217 bp. The overall base composition of the major coding strand was A 41.6%, T 43.1%, C 7.1%, and G 8.2%, with a high A+T bias of 84.7%.

**Figure 1.**
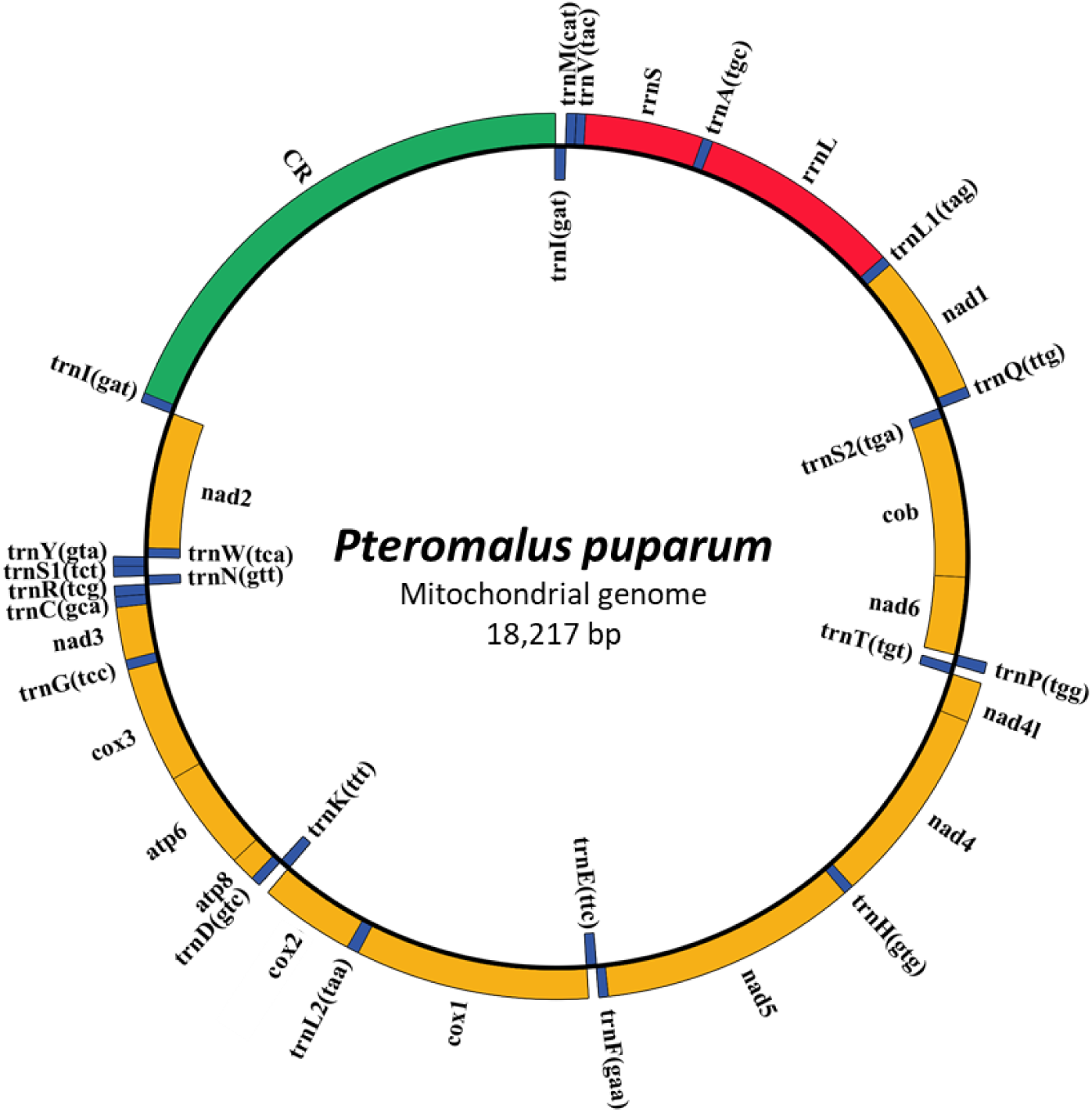
Structure of *Pteromalus puparum* mitochondrial genome. Yellow indicates mitochondrial protein-coding genes. Red indicates mitochondrial ribosomal RNAs. Blue indicates mitochondrial tRNAs. Green indicates mitochondrial control region, which is also known as mitochondrial A+T-rich region, D-loop region or major non-coding region. Coding genes outside the circle are encoded by the major coding strand, and inside genes are encoded by the minor strand.

Mitochondrial genes were annotated using the MITOS web server (Bernt *et al.,* 2013). All 37 typical mitochondrial genes were detected (Figure 1, Table 1). In addition to the standard set of genes, an extra identical isoleucine tRNA (trnI, mitochondrial gene abbreviations are taken from Cameron (2014)) was also found (Figure 1, Table 1). These two identical trnI tRNAs are on opposite ends with each adjacent to the control region. One is coded on the major coding strand, the other is coded on the minor strand. This additional mitochondrial gene is an example of recent gene duplication in mitochondrial genome. Although gene duplication is less common in mitochondrial genomes, it has also been observed in several other insects (Beckenbach 2011; Junqueira *et al.,* 2004; Kim *et al.,* 2006; Kocher *et al.,* 2014), and proposed as ongoing process of gene order rearrangement (Beckenbach 2011; Beckenbach 2012; Moritz *et* al., 1987).

**Table 1.** Summary of *Pteromalus puparum* mitochondrial genes.

| Name | Strand | Start | Stop | Length<sup>a</sup> | *Nasonia vitripennis*(AsymC) | *Philosophy pilosa* |
| --- | --- | --- | --- | --- | --- | --- |
| trnI(gat) | – |  |  | 69 | not found | 67 |
| trnM(cat) | + |  |  | 66 | not found | not found |
| trnV(tac) | + |  |  | 66 | not found | not found |
| rrnS | + |  |  | 778 | 722 | incomplete |
| trnA(tgc) | + |  |  | 65 | 64 | 65 |
| rrnL | + |  |  | 1362 | 1331 | 1309 |
| trnL1(tag) | + |  |  | 68 | 67 | 63 |
| nad1 | + | ATA | TAG | 933 | 933 | 919 |
| trnQ(ttg) | + |  |  | 72 | not found | 70 |
| trnS2(tga) | – |  |  | 68 | not found | 66 |
| cob | – | ATG | TAA | 1140 | 1137 | 1140 |
| nad6 | – | ATG | TAA | 552 | 546 | 564 |
| trnP(tgg) | + |  |  | 66 | 65 | 66 |
| trnT(tgt) | – |  |  | 64 | 65 | 68 |
| nad4l | + | ATT | TAA | 264 | 291 | 285 |
| nad4 | + | ATG | T<sup>b</sup> | 1332 | 1338 | 1341 |
| trnH(gtg) | + |  |  | 63 | 65 | 67 |
| nad5 | + | ATA | TAA | 1680 | 1686 | 1666 |
| trnF(gaa) | + |  |  | 64 | 65 | 66 |
| trnE(ttc) | – |  |  | 65 | 71<sup>c</sup> | 69 |
| cox1 | + | ATA | TAA | 1539 | 1557 | 1545 |
| trnL2(taa) | + |  |  | 66 | 68 | 66 |
| cox2 | + | ATA | TAA | 651 | 666 | 673 |
| trnK(ttt) | – |  |  | 72 | 74 | 69 |
| trnD(gtc) | + |  |  | 68 | 65 | 65 |
| atp8 | + | ATT | TAA | 159 | 156 | 165 |
| atp6 | + | ATG | T<sup>b</sup> | 669 | 672 | 675 |
| cox3 | + | ATA | TAA | 774 | 783 | incomplete |
| trnG(tcc) | + |  |  | 67 | 68 | not found |
| nad3 | + | ATA | TAA | 339 | 349 | 334 |
| trnC(gca) | + |  |  | 66 | not found | 65 |
| trnR(tcg) | + |  |  | 70 | not found | not found |
| trnN(gtt) | – |  |  | 66 | 67 | not found |
| trnS1(tct) | + |  |  | 60 | not found | 62<sup>c</sup> |
| trnY(gta) | + |  |  | 66 | 68 | 66 |
| trnW(tca) | – |  |  | 68 | 67<sup>c</sup> | 66 |
| nad2 | – | ATA | TAA | 969 | 975 | 969 |
| trnI(gat) | + |  |  | 69 | not found | 67 |

Genes in *P. puparum* mitochondrial genome were annotated using MITOS, followed by manually adjustment for protein-coding genes. Statistics were retrieved from Oliveira *et al.* (2008) and Xiao *et al.* (2011) for genes in *N.* *vitripennis* and *P. pilosa* mitochondrial genomes, respectively. + indicates the major coding strand; – indicates the
minor strand.
<sup>a</sup> The length of protein-coding genes is given excluding the stop codon.
<sup>b</sup> Stop codon is single T, and will be completed by posttranscriptional polyadenylation.
<sup>c</sup> This tRNA was annotated by MITOS in this paper.

All 24 annotated tRNAs show canonical cloverleaf secondary structures, except trnS1 (Figure S2). Mitochondrial trnS1 lacks the dihydrouridine (DHU) arm that is common in most metazoans (Cameron 2014; Juhling *et al.,* 2012). The start and end positions of 13 protein-coding genes were manually checked, and finally adjusted in 9 of 13 genes (Table 1). The lengths of these protein-coding genes were compared and found similar to those of the pteromalid *N. vitripennis* and *P. pilosa* (Table 1).

In addition to conserved coding genes, the 3,308 bp control region was also investigated. This control region is highly AT-rich. The overall base composition of the major coding strand is A 42.4%, T 43.5%, C 7.0%, and G 7.1%, with a high A+T bias of 85.9%. Eight tandem repeats followed by 6 reverse repeats were detected in this control region by Windotter (Figure S3). The consensus sequence is 206 bp (Figure 2). Of these 14 repeats, repeat 2, 3, 5, 6 and 7 are identical to the census sequence (Figure 2). Mismatches and/or indels were found in the other repeats. Repeat 8 (136 bp) and 9 (163 bp) are incomplete compared to the consensus sequence (206 bp) (Figure 2). This long and complex control region probably explains prior failures to amplify and sequence the complete mitochondrial genomes in most chalcidoids (Nedoluzhko *et al.,* 2016; Oliveira *et al.,* 2008; Su *et al.,* 2016; Xiao *et al.,* 2011; Xiao *et al.,* 2012).

**Figure 2.**
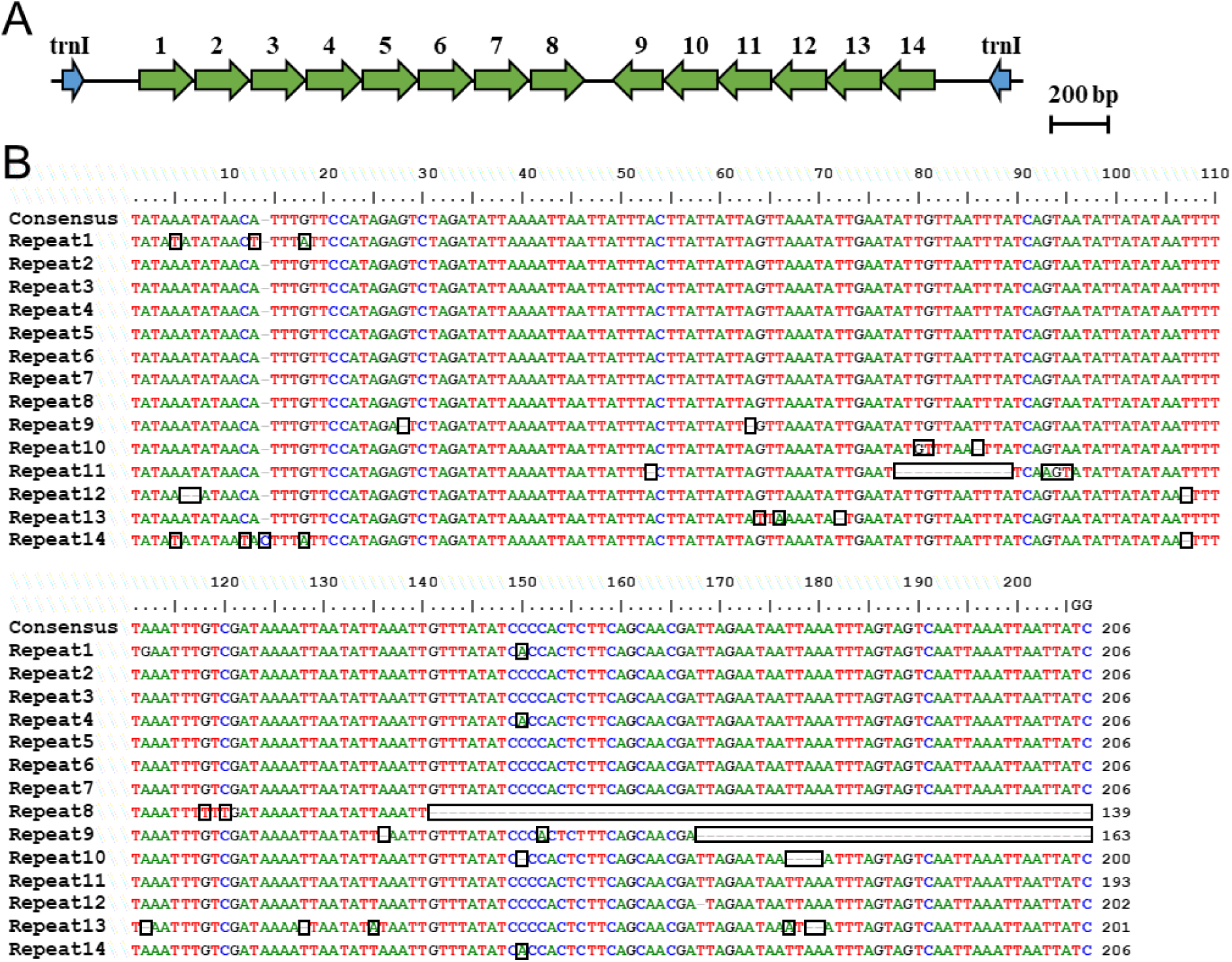
Repeats in control region of *P. puparum* mitochondrial genome. **(A)** Shown is 8 repeats followed by 6 reversed repeats identified in control region of *P. puparum* mitochondrial genome. **(B)** Also shown is the alignment of these 14 repeats. The consensus sequence of repeats is 206 bp. Differences to the consensus sequence are highlighted by rectangles.

### 2, Frequent gene order rearrangement in Chalcidoidea

First, we compared mitochondrial gene orders of *P. puparum* to *N. vitripennis*, its closest taxon with available mitochondrial genome information. Despite their close relationship within the same subfamily Pteromalinae, we observed gene order rearrangement that are long range transpositions of trnN and “trnY-trnW-nad2” block, which contains a protein-coding gene (Figure 3). This indicates that Pteromalinae belongs to one of the groups with high rates of gene order rearrangement.

**Figure 3.**
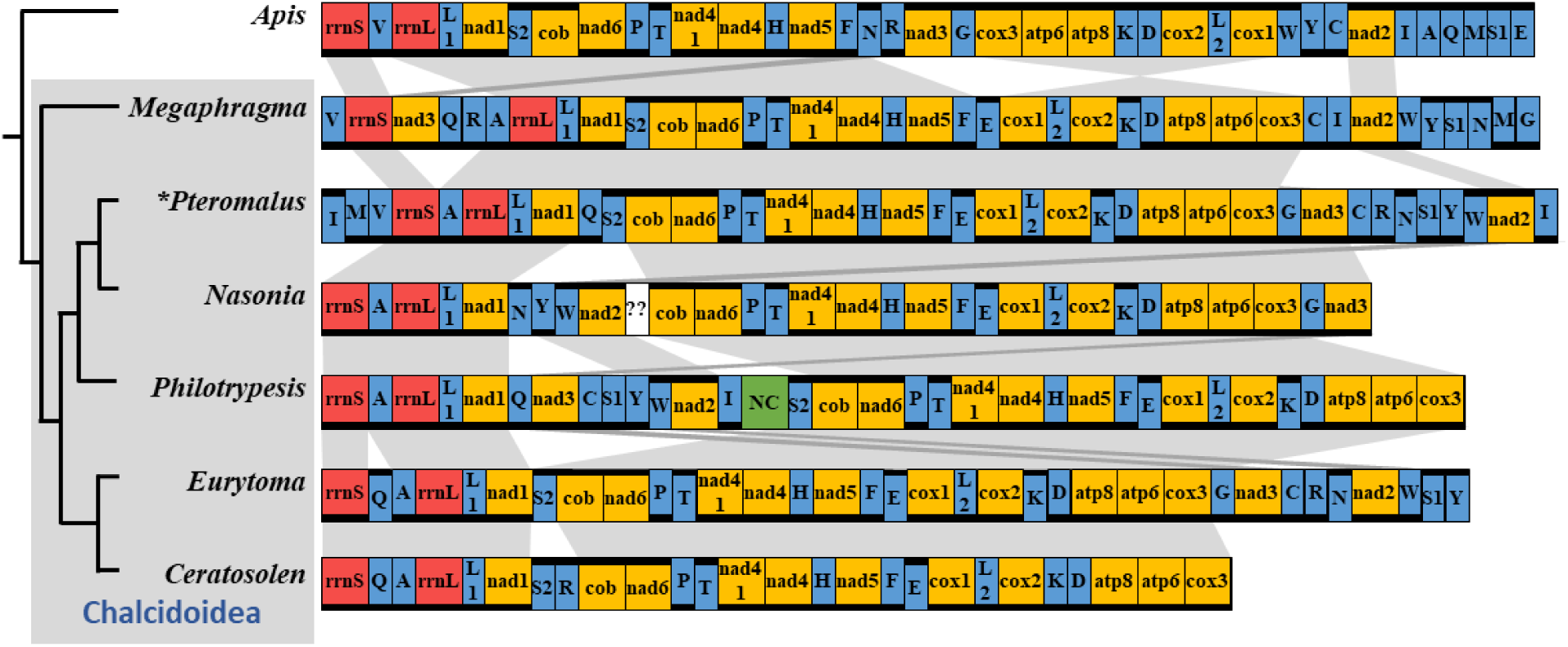
Rearrangement of mitochondrial gene order in *P. puparum*. Yellow indicates mitochondrial protein-coding genes. Red indicates mitochondrial ribosomal RNAs. Blue indicates mitochondrial tRNAs. tRNA genes are indicated by the single527 letter abbreviations for their corresponding amino acid. Green indicates mitochondrial non-coding region identified in *Philotrypesis* mitochondrial genome. Genes encoded by the major strand, which encodes mitochondrial ribosomal RNAs, are labeled with black bold lines under the boxes. Genes encoded by the opposite minor strand are labeled with black bold lines on the top of boxes. Large blocks and protein-coding genes were aligned by gray shadows. Asterisk indicates that complete circular mitochondrial genome is available. High rate of mitochondrial gene order rearrangement was observed in Chalcidoidea.

We further investigated mitochondrial gene order rearrangements in other Chalcidoidea. To get available mitochondrial genomes in Chalcidoidea, blastn of *P. puparum* cox1 was conducted on NCBI by setting organism limited to Chalcidoidea (2018-March). The longest mitochondrial genome records were chose when there were multiple hits in the same genus. Four additional partial mitochondrial genomes were retrieved. They are mitochondrial genomes from *Philotrypesis pilosa* (Pteromalidae), *C. solmsi* (Agaonidae), *Eurytoma* sp. TJS-2016 (Eurytomidae) and *Megaphragma amalphitanum* (Trichogrammatidae).

Among all these 6 chalcidoids, no two species share the same mitochondrial gene order (Figure 3). The smallest difference was found between truncated *C. solmsi* and partial *Eurytoma* sp. TJS-2016 mitochondrial genome, where trnQ and trnK were locally reversed (i.e. moved to the opposite strand at same position), and trnR were long-range transposed. In comparison to ancestral insect mitochondrial genome, a striking gene rearrangement is reversion of a big block “cox1-trnL2-cox2-trnK-trnD-atp8-atp6-cox3” shared in all 6 chalcidoids (Oliveira *et al.,* 2008; Xiao *et al.,* 2011). Another conversed block “trnA-rrnL-trnL1-nad1” were shared in chalcidoids compared to other insects. For other mitochondrial genes, multiple rearrangement events were observed within Chalcidoidea for both mitochondrial tRNAs and protein-coding genes (Figure 3). Nad2 and nad3 are ‘hot spots’ for mitochondrial gene rearrangement within Chalcidoidea. Interestingly, a non-coding region was found inserted between “nad2-trnI” and “trnS2-cob” in *P. pilosa* (Xiao *et al.,* 2011). Corresponding regions in *Nasonia* were not able to be amplified and sequenced (Oliveira *et al.,* 2008). Unexpected non-coding region may explain the failure of joining two separate mitochondrial PCR fragments in *Nasonia* (Oliveira *et al.,* 2008).

### 3, Protein substitution rate comparison in Chalcidoidea

Previous studies reported strikingly high substation rates of mitochondrial genomes in Chalcidoidea wasps (Oliveira *et al.,* 2008; Xiao *et al.,* 2011; Xiao *et al.,* 2012). For example, mitochondrial synonymous substitution rate between *N. giraulti* and *N. longicornis* is 45.24%, which is 40 times higher than nuclear synonymous substitution rate (1.12 %) between these two species (Oliveira *et al.,* 2008; Raychoudhury *et al.,* 2009a). We first estimated synonymous substitution rates between *P. puparum* and *N. vitripennis* mitochondrial genes, using Yang and Nielsen (2000) method for correcting multiple substitutions at the same site. Among 13 mitochondrial protein-coding genes, the smallest synonymous substitution is 610 % for cox1, and the average is 8130 % (Table S2), which is much higher than reported synonymous rate (30.8 %) from a nuclear gene calreticulin (Yan *et al.,* 2016). Thus, synonymous substitution rates between *P. puparum* and *N. vitripennis* mitochondrial genomes are very fast and highly saturated.

We then investigated protein substitution rates for Chalcidoidea mitochondrial genomes. First, protein substitution rates were estimated using concatenated mitochondrial proteins (Figure 4A). As information of nad2 and nad3 were not available in *C. solmsi* (Xiao *et al.,* 2012), the remaining 11 mitochondrial proteins were concatenated for analysis. Considerable rate variation was observed among Chalcidoidea mitochondrial genomes. *Ceratosolen solmsi* (Agaonidae) shows the longest protein distances to Chalcidoidea common ancestor, and *M. amalphitanum* (Trichogrammatidae) shows the shortest (Figure 4A). Protein distance to the Chalcidoidea common ancestor is ∼6.5 times higher in *C. solmsi* than that in *M. amalphitanum*. For the remaining four chalcidoids, protein distances to the Chalcidoidea common ancestor are similar, although *P. puparum* is slightly shorter than the other three chalcidoids (Figure 4A).

**Figure 4.**
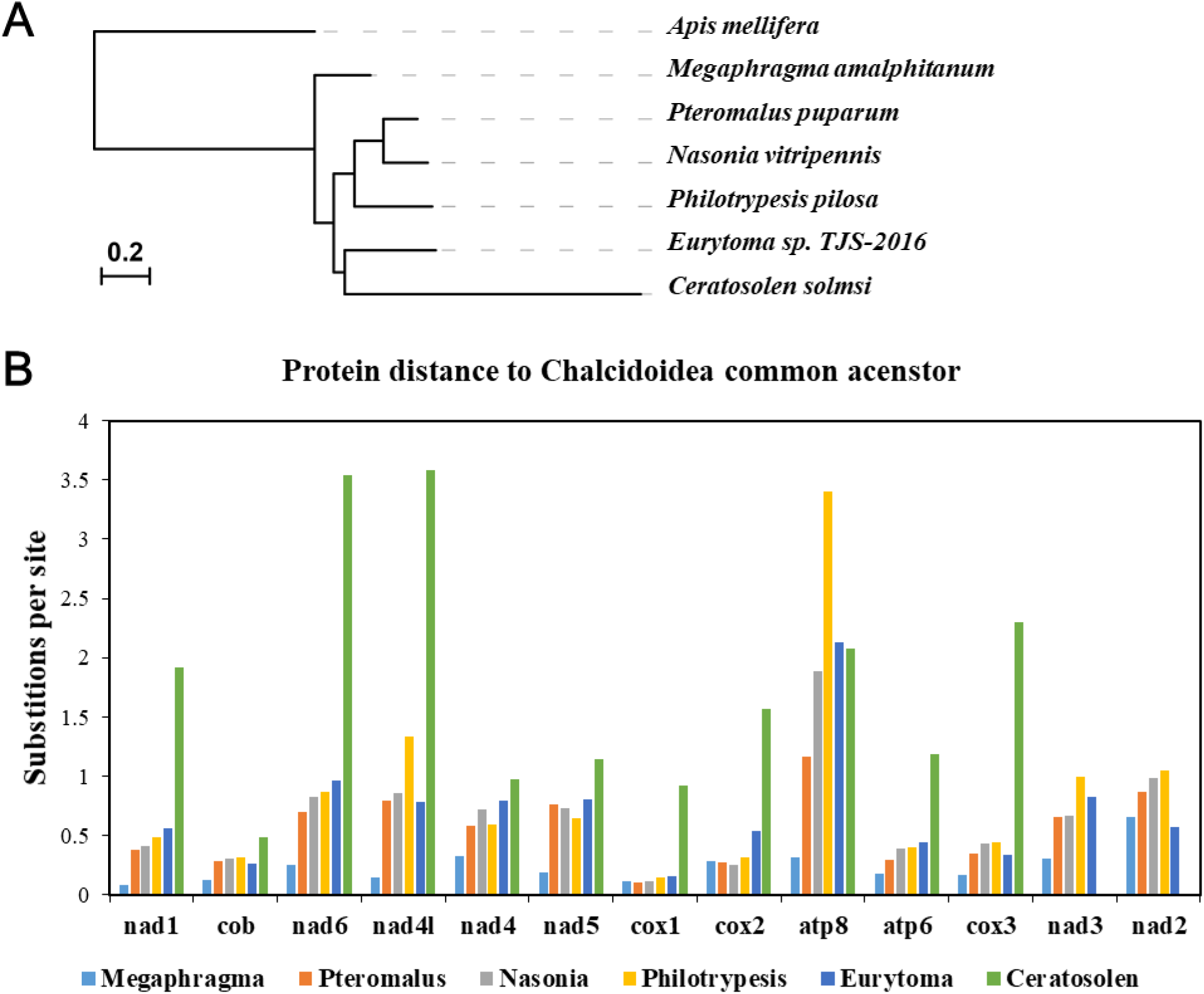
Comparison of mitochondrial protein distances to Chalcidoidea common ancestor. **(A)** Protein substitutions were estimated using concatenated 11 mitochondrial proteins. Information of nad2 and nad3 were not available in *Ceratosolen solmsi*. **(B)** Distances to Chalcidoidea common ancestor for 13 individual mitochondrial proteins. Mitochondrial protein substitution rates of *P. puparum* are comparable to those of *Nasonia vitripennis*. *Ceratosolen solmsi* show higher protein substitution rates than other chalcidoids for all mitochondrial proteins, except atp8.

To test whether a similar tendency is observed across the entire mitochondrial genome, individual mitochondrial protein analyses were also conducted. For 11 available mitochondrial proteins in *C. solmsi, C. solmsi* shows the highest protein substitution rates among 6 chalcidoids, except atp8 (Figure 4B). In contrast, *M. amalphitanum* shows the slowest protein substitution rates in 10 of 13 mitochondrial proteins. Distance to the Chalcidoidea common ancestor for *P. puparum* is significantly longer than that for *M. amalphitanum*, and significantly shorter than those of other chalcidoids (Figure 4B; Wilcoxon signed rank test followed by Benjamini & Hochberg multiple correction, *p* < 0.05 for each case). However, the rates are similar in the family Pteromalidae, which includes *P. puparum, N. vitripennis* and *P. pilosa*, indicating high protein substitution rate also in the *P. puparum* mitochondrial genome.

### 4, NUMTs identified in *Pteromalus puparum* genome

We investigated nuclear insertions of mitochondrial DNA (NUMTs) in the *P. puparum* genome, using the same method reported for *N. vitripennis* and *A. mellifera* (Behura 2007; Viljakainen *et al.,* 2010). Totally, 24 NUMTs summing up to 9,989 bp were found scattered over 22 genomic scaffolds in *P. puparum* (Table S3). Thus, total length of NUMTs in the *P. puparum* genome is less than a quarter of that reported in *N. vitripennis* (42,972 bp) (Viljakainen *et al.,* 2010), and much less than total NUMTs length in *A. mellifera* (over 230 kb) (Behura 2007). Considering that the *Nasonia* mitochondrial genome is only partially sequenced and NUMTs of the control region were therefore not detectable, the difference of total NUMTs length between *P. puparum* and *N. vitripennis* is even more pronounced. Only 11 of 24 NUMTs in *P. puparum* originated from mitochondrial coding regions (Table S3). These NUMTs of coding regions summed up to 3,617 bp, only around one-twelfth of that found in *N. vitripennis*.

To rule out the possibility that total NUMTs length difference is caused by difference of methods, e.g. blast version or subtle setting differences, we also undertook NUMTs identification in *N. vitripennis* and *C. solmsi* genomes. Our method retrieved all regions of previously reported in *N. vitripennis* genome. Using *C. solmsi* partial mitochondrial genome which only contain 11 of 13 mitochondrial proteins, we found 176 NUMTs summing up to 53,962 bp in the *C. solmsi* genome (Xiao *et al.,* 2013), which is around fifteen-fold greater than *P. puparum*. These results indicate that the shorter total NUMTs length detected in *P. puparum* is not caused by a method difference.

It has been reported that NUMT length is positively correlated with genome size, suggesting potential roles of non-coding DNA gain and loss in NUMT accumulation (Hazkani-Covo *et al.,* 2010; Song *et al.,* 2013). However, genome size can’t explain fewer NUMTs in *P. puparum*, as assembled genome size of *P. puparum* is larger than those of *N. vitripennis, A. mellifera* and *C. solmsi* (Weinstock *et al.,* 2006; Werren *et* al., 2010; Xiao *et al.,* 2013). In addition, fewer detected NUMTs in *P. puparum* is not caused by faster evolutionary rate of *P. puparum* mitochondrial genome, as slower mitochondrial protein substitution rates were shown in *P. puparum* than *N. vitripennis* and *C. solmsi* (Figure 4). Genome assembling difference is also unlikely to cause less NUMTs in *P. puparum* genome. Because no filtering of mitochondrial sequences had been applied to *P. puparum* sequencing and assembling, and mitochondrial genome sequence Scaffold225127 was detected in *P. puparum* genome assemblies. Using combined PacBio and Illumina sequencing, assembly quality of *P. puparum* genome is better than that of *N. vitripennis* (Werren *et al.,* 2010). It is also less likely to fail in assembling NUMTs in *P. puparum*.

To get more comprehensive understandings of NUMTs in *P. puparum*, phylogenetic analyses were conducted. For NUMTs of protein-coding genes, 4 were found longer than 100 bp. They are two NUMTs containing cox1, one cox2 and one nad4. We named them *Ppup*NUMT1–4, respectively. Maximum likelihood phylogenetic trees were constructed for these NUMTs with mitochondrial protein-coding sequences. NUMTs were clustered with mitochondrial genes from the same species (Figure 5). These indicate frequent dynamics of NUMTs transferring and degrading. Strikingly, branches leading to *Ppup*NUMT2 and *Ppup*NUMT3 are extremely long, compared to branches leading to *P. puparum* mitochondrial genes and NUMTs in *N. vitripennis* (Figure 5). These results suggest that NUMTs in *P. puparum* evolve faster after transferring into nuclear genome than that in *N. vitripennis*, and are more likely to degrade to the point where they are undetectable by blast. We speculate that a fast degrading rate of NUMTs is one possible reason for fewer detected NUMTs in *P. puparum* genome. However, the sampling size here is too small for an efficient statistical test. In addition, other factors, such as lower integration rate into the nuclear genome or higher deletion rate of NUMTs, can also contribute to fewer detected NUMTs in *P. puparum*. Further investigations are needed to compare rates of insertion, degradation or deletion of NUMTs in more chalcidoid genomes.

**Figure 5.**
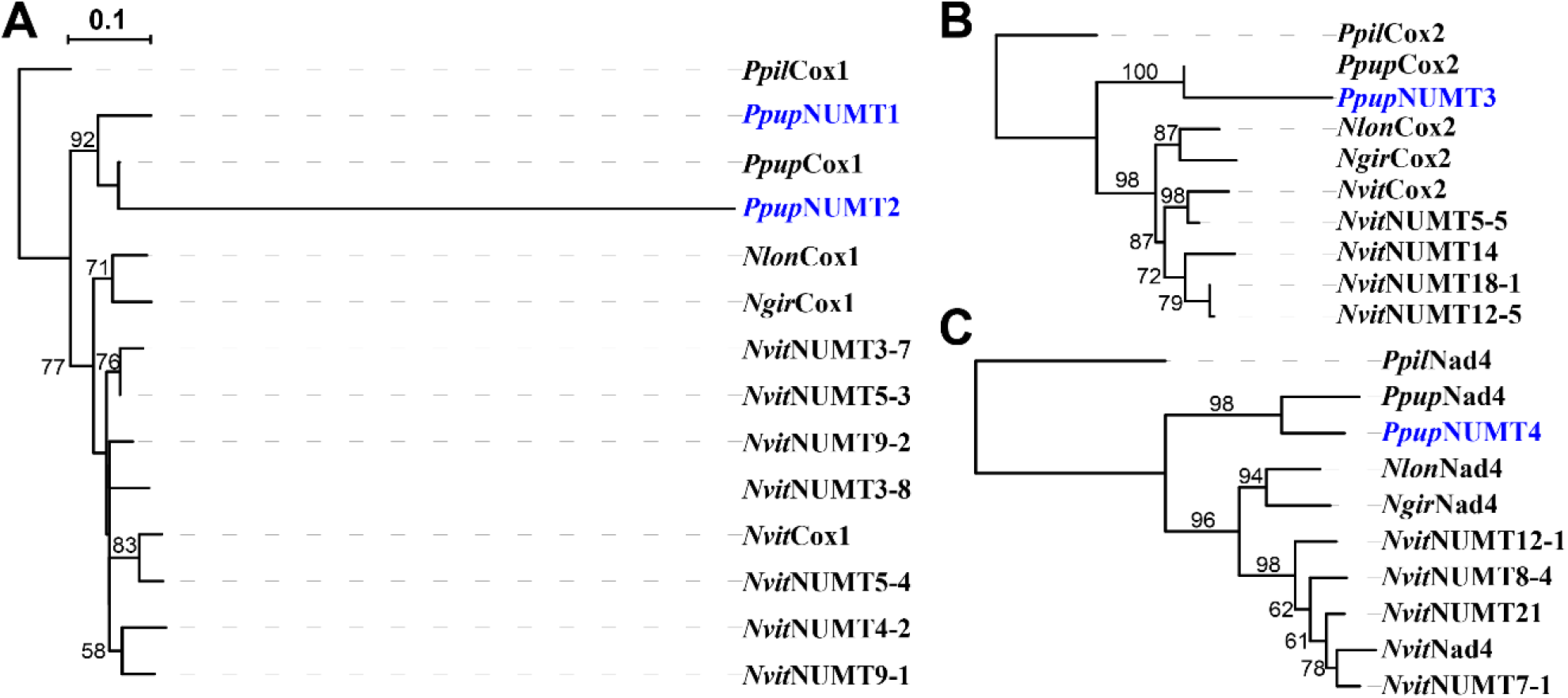
NUMTs of mitochondrial protein-coding genes in *P. puparum* genome. Phylogenetic analysis of NUMTs of **(A)** Cox1, **(B)** Cox2 and **(C)** Nad4. Mitochondrial protein-coding genes from *P. pilosa* were used as outgroups. Numbers are bootstrap supports for the nodes when higher than 50. NUMTs in *P. puparum* genome were labelled blue. The regions used were Scaffold115:44107-44450 for *Ppup*NUMT1, Scaffold625:135190-135914 for *Ppup*NUMT2, Scaffold263595:2668-2931 for *Ppup*NUMT3 and Scaffold22:1352012-1352494 for *Ppup*NUMT4. *Ppil*: *P. pilosa*; *Ppup*: *P. puparum*; *Nvit*: *N. vitripennis*; *Ngir*: *N. giraulti*; *Nlon*: *N. longicornis*.

## Conclusions

Our study reports a circular mitochondrial genome from an endoparasitoid wasp, *P. puparum*. Comparative analyses with other chalcidoids reveal frequent gene order rearrangement and variable protein evolutionary rates in Chalcidoidea mitochondrial genomes. Our study also reveals a dramatic difference of NUMTs abundance between *P. puparum* and its relatives. Phylogenetic analyses suggest faster degradation rate of NUMTs in *P. puparum*, which can lead to fewer NUMTs. Overall, our study provides more information and understanding about evolution of both mitochondrial genomes and NUMTs in Chalcidoidea.

## Methods

### 1, Mitochondrial genome assembly

To retrieve mitochondrial genome from data of *P. puparum* genome sequencing projects, *N. vitripennis* (AsymC) mitochondrial sequences EU746609.1 and EU746613.1 were used as queries for blastn against *P*. *puparum* genome assembly. The best hit was Scaffold225127 as potential *P. puparum* mitochondrial genome sequence. Illumina and PacBio reads were then aligned to Scaffold225127 using BWA v0.7.4 (Li & Durbin 2009). Combined Illumina and PacBio reads, the mitochondrial genome was assembled and circulated. The assembly was completed by Nextomics Biosciences Co., Ltd. (China). The deposited GenBank accession is MH051556. To evaluate assembly of control region, the circular mitochondrial genome was cut into linear at middle of coding region, which is corresponding position 10,000 in GenBank sequence MH051556. PacBio reads were then aligned to the cut mitochondrial genome using BWA (Li & Durbin 2009). Supporting PacBio reads were viewed in IGV 2.3.91 (Robinson *et al.,* 2011).

### 2, Retrieving other mitochondrial genomes in Chalcidoidea

To retrieve available Chalcidoidea mitochondrial genomes, blastn of *P. puparum* cox1 sequence was conducted against NCBI nr database by limiting organism to “Chalcidoidea”. “Max target sequences” was set as 20,000. In this paper, the sequences used are mitochondrial genomes of *N. vitripennis* (GenBank EU746613.1, EU746609.1), *N. giraulti* (EU746614.1, EU746611.1), *N. longicornis* (EU746616.1, EU746612.1), *P. pilosa* (JF808723.1), *C. solmsi* (JF816396.1), *Eurytoma* sp. TJS-2016 (KX066374.1) and *M. amalphitanum* (KT373787.1).

### 3, Mitochondrial genome annotation

Genes from the *P. puparum* mitochondrial genome were annotated using MITOS web server (Bernt *et al.,* 2013). Start and stop positions of protein coding genes were manually adjusted. Tandem repeats in control region were discovered by Windotter (Sonnhammer & Durbin 1995) and identified by online Tandem repeats finder (Benson 1999) https://tandem.bu.edu/trf/trf.html. The repeats were aligned by MUSLE v3.8.31 (Edgar 2004), and visualized in BioEdit v7.2.5 (Hall 1999).

### 4, Evolutionary rate estimation

Pairwise synonymous and non-synonymous substitution rates were estimated using yn00 in PAML v4.8 (Yang 2007; Yang & Nielsen 2000). For protein substitution rate estimation, mitochondrial protein sequences were aligned by Mafft v7.123b using L-INS-i mode (Katoh & Standley 2013), followed by filtering using trimAI v1.4.1 (Capella-Gutierrez *et al.,* 2009). The alignments were concatenated using AMAS (Borowiec 2016). Protein substitution rates were estimated based given tree using RAxML v 8.0.20 by setting substitution model as “PROTGAMMAAUTO” (Stamatakis 2014). The tree topology is inferred from Peters *et al.* (2018). Distances to Chalcidoidea common ancestor were extracted using Newick Utilities v1.6 (Junier & Zdobnov 2010). Statistical analyses were performed in R v3.2.0. Wilcoxon signed rank test was used for significance test of differences, followed by Benjamini & Hochberg multiple correction.

### 5, NUMTs identification and phylogenetic analysis

To identify NUMTs, *P. puparum* mitochondrial genome sequence (GenBank MH051556) was used as query for blastn against *P. puparum* genome (excluding mitochondrial Scaffold225127). The search was done using blast v2.6.0+ with E-value < 0.0001. Initially, 157 hits were detected in *P. puparum* using blastn with E-value < 1e^-4^. Several hits share overlapping, especially for hits of repeats from the mitochondrial control region. These overlapped hits were manually joined. For mitochondrial protein coding genes, their NUMTs longer than 100 bp were used for phylogenetic analysis. Blastn of individual mitochondrial protein coding sequences were conducted against nuclear genome to determine the corresponding regions of NUMTs. Mitochondrial protein coding genes and their NUMTs were aligned using built-in MUSLE in MEGA v7 (Kumar *et al.,* 2016). Phylogenetic trees were constructed in MEGA v7 using maximum likelihood method based on GTR+G model. The maximum likelihood trees were run for 1,000 bootstrap replicates. The trees were visualized using iTOL (Letunic & Bork 2016).

## Acknowledgements

This research was supported by Major International (Regional) Joint Research Project of NSFC (Grant no. 31620103915 to GY and JHW), National Natural Science Foundation of China (Grant no. 31472038 to GY, no. 31701843 to ZY), US National Science Foundation (DEB-1257053 and IOS-1456233) and Nathaniel & Helen Wisch Chair to JHW and China Postdoctoral Science Foundation (2016M601947) to ZY. In addition, it is also supported by the Program for Chinese Outstanding Talents in Agricultural Scientific Research of Ministry of Agriculture and Rural Affairs, and the Program for Chinese Innovation Team in Key Areas of Science and Technology of Ministry of Science and Technology (2016RA4008).

## Author Contributions

GY, JHW and XC conceived and designed the research; YT assembled the mitochondrial genome; FQ and FW collected the online data; ZY performed analyses; GY, JHW, FQ, XC and ZY interpreted the results; GY, JHW and ZY wrote and revised the manuscript.

